# Spontaneous oligomerization of BAK/BAX is suppressed by hetero-dimerization with MCL-1

**DOI:** 10.1101/756874

**Authors:** Basile I. M. Wicky, Kallol Gupta, Tristan O. C. Kwan, Carol V. Robinson, Jane Clarke

**Author notes:** Corresponding authors (C.V.R.; J.C.).

## Abstract

BCL-2 proteins control the intrinsic pathway of programmed cell death. Composed of anti- and pro-apoptotic members, their network of interactions forms a molecular switch that controls mitochondrial outer-membrane permeability. Apoptotic stimulation leads to BAK/BAX oligomerization and pore formation, yet the molecular details of this pivotal step remain poorly understood, and controversy persists regarding the activation mechanism. Here we use native mass spectrometry and kinetics to show that the homo-oligomerization of BAK and BAX is spontaneous in hydrophobic environments. This process is abrogated by hetero-dimerization of both BAK and BAX with the anti-apoptotic BCL-2 protein MCL-1. Pro-apoptotic BH3-only proteins disrupt these hetero-dimers by binding competitively to MCL-1, releasing BAK/BAX for homo-oligomerization. Thus, we infer that their oligomeric states are thermodynamically favored at the membrane. Our approach provides the framework for future quantitative biophysical characterizations of the BCL-2 network, and advances our molecular understanding of apoptosis.

---

A number of models have been proposed to explain BAK/BAX activation and pore-formation (Chipuk & Green, 2008; Czabotar, Lessene, Strasser, & Adams, 2014; Kale, Osterlund, & Andrews, 2017; Peña-Blanco & García-Sáez, 2018). On one hand, ‘direct’ activation models stipulate that the trigger is a physical interaction between BH3-only proteins and BAK/BAX (Kim et al., 2006; Kuwana et al., 2005; Letai et al., 2002). Alternatively, ‘indirect’ activation models posit that BH3-only proteins do not engage BAK/BAX, but instead neutralize the anti-apoptotic BCL-2 proteins that prevent their oligomerization under non-apoptotic conditions (Uren et al., 2007; Willis et al., 2007). To study the BCL-2 network, we selected a minimal set of proteins encompassing all functional sub-classes. Both BAK and BAX (pore-forming) were investigated, MCL-1 was chosen as the model anti-apoptotic protein, and pro-apoptotic BH3-only proteins (PUMA and BID) were studied as 35-residue peptides (more details about the BCL-2 family can be found in Extended Data Fig. 1). We used native MS and biophysical techniques to probe the interactions between the components of the system, elucidate oligomeric stoichiometries, and investigate thermodynamic and kinetic aspects of these processes.

First, we analyzed interactions between the components under simple aqueous buffer conditions. In line with previous reports, the affinities between BH3 motifs and MCL-1 were all found to be tight (low to sub-nM, *cf.* Table 1) (Dahal, Kwan, Hollins, & Clarke, 2018; Kong et al., 2018; Ku, Liang, Jung, & Oh, 2011). Importantly, all bimolecular association rate constants are fast (10^6^–10^7^ M^-1^ s^-1^), and differences in affinities are almost entirely due to changes in lifetimes of the complexes (*t*_1/2_ ≈ 1 s to 20 min). Certain BH3-only proteins (termed ‘activator’, *e.g.* PUMA, BID) have been reported to directly engage BAK/BAX, triggering their oligomerization (Dai, Pang, Ramirez-Alvarado, & Kaufmann, 2014; Gavathiotis et al., 2008; Kuwana et al., 2005; Letai et al., 2002; Moldoveanu et al., 2013). Given the importance of these postulated interactions for the ‘direct’ activation model, the paucity of binding data available is surprising. We measured binding isotherms between BH3 peptides of PUMA and BID with BAK (Table 1 and Extended Data Fig. 2a). Surprisingly, these interactions were found to be orders of magnitudes weaker (high μM range) than the binding of the same peptides to MCL-1 (Table 1) (Dahal et al., 2018; Kong et al., 2018; Ku et al., 2011), suggesting that these binding events are irrelevant under physiological concentrations of these proteins (*e.g.* the concentration of BAX has been reported in the range of 2.2–170 nM (Eskes et al., 1998; Polster, Basañez, Young, Suzuki, & Fiskum, 2018)). This sharp dichotomy is striking given the structural homology between BAK/BAX and anti-apoptotic BCL-2 proteins (Extended Data Fig. 3), and might suggest that BAK:BH3 interactions were selected *against* over the course of evolution. We note that these very weak affinities are solely the consequences of shortened lifetimes of the bound states (*t*_1/2_ ≈ 0.5–3 ms), since association rate constants to BAK and MCL-1 are similar (Table 1). While, these kinetic profiles might appear to support the ‘hit-and-run’ mechanism (Chipuk & Green, 2008; Dai et al., 2011), we observed a complete absence of oligomerization by size-exclusion chromatography, even when 10-fold excesses of BH3 peptides were employed (Extended Data Fig. 4). Thus, we found no evidence to suggest that these BH3-only proteins were ‘activators’ capable of triggering the oligomerization of BAK/BAX on their own. Surprisingly, we also observed a lack of complex formation between BAK and MCL-1 (Extended Data Fig. 2c). Since the binding between BAK_BH3_ and MCL-1 is very tight (*K*_d_ = 0.08 nM, *cf.* Table 1), we infer that the BH3 motif is inaccessible in the context of folded monomeric BAK, and that the energetic cost of conformational changes and/or partial unfolding cannot be offset by the interaction alone. Taken together, these results show that ‘standard’ biochemical conditions do not recapitulate the behavior of the BCL-2 network observed in cells.

**Table 1.**
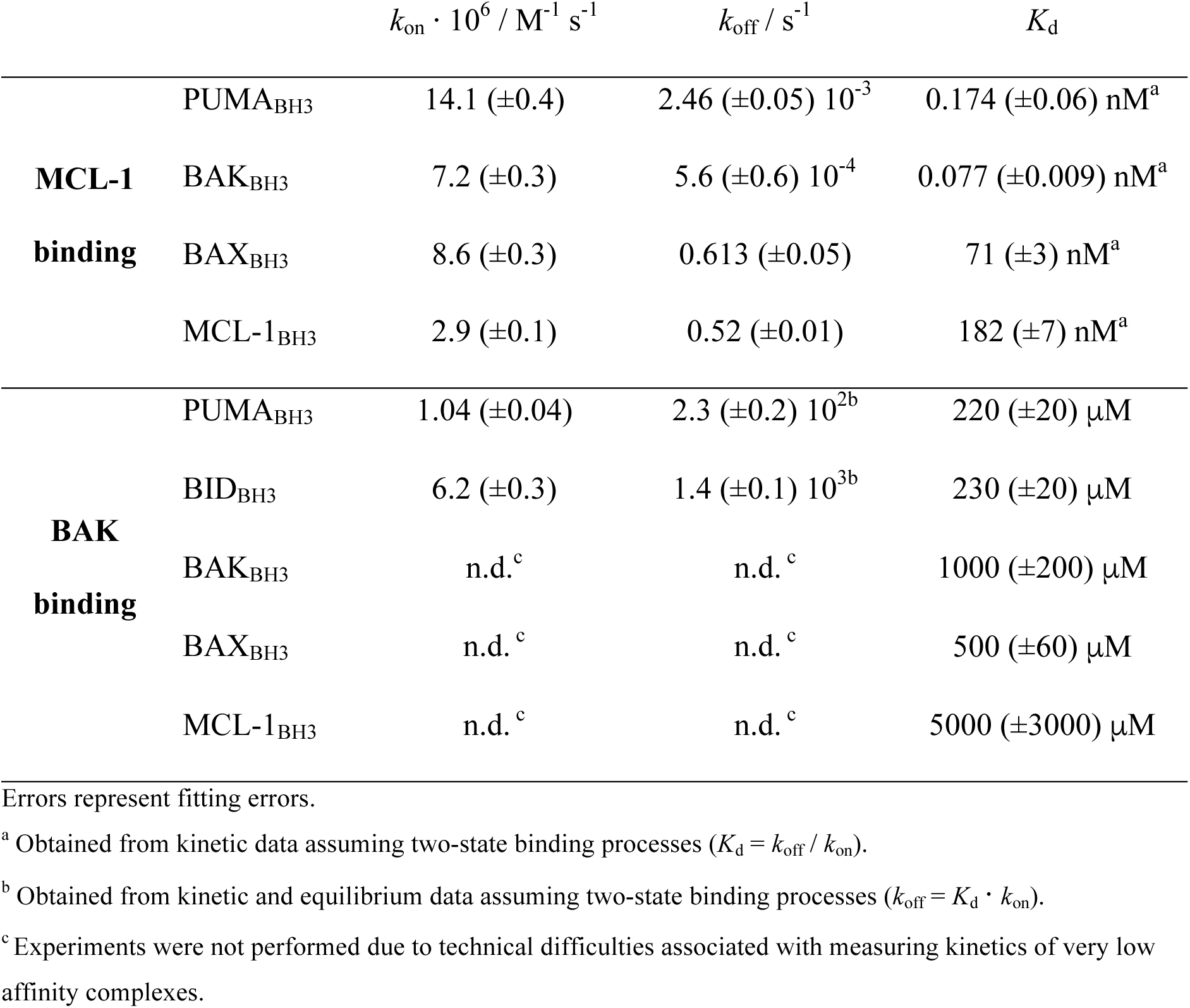
Kinetic and thermodynamic parameters for the binding of BH3 peptides to BAK and MCL-1 in buffer without detergent.

Some BCL-2 proteins are cytosolic, but many are localized at the mitochondrial outer-membrane *via* single-pass C-terminal helices, including BAK and BAX (Schellenberg et al., 2013). We found that under membrane mimetic detergent micelle conditions, both proteins spontaneously formed higher-order oligomers, as revealed by size-exclusion chromatography and chemical cross-linking (Fig. 1a–c). In contrast, the anti-apoptotic MCL-1 remained monomeric (Extended Data Fig. 5c), denoting its distinct biological function. These oligomeric structures were folded, as confirmed by CD spectroscopy, although a slight loss of helicity and change in tryptophan fluorescence were clear indictors of structural rearrangements (Extended Data Fig. 6).

**Fig. 1.**
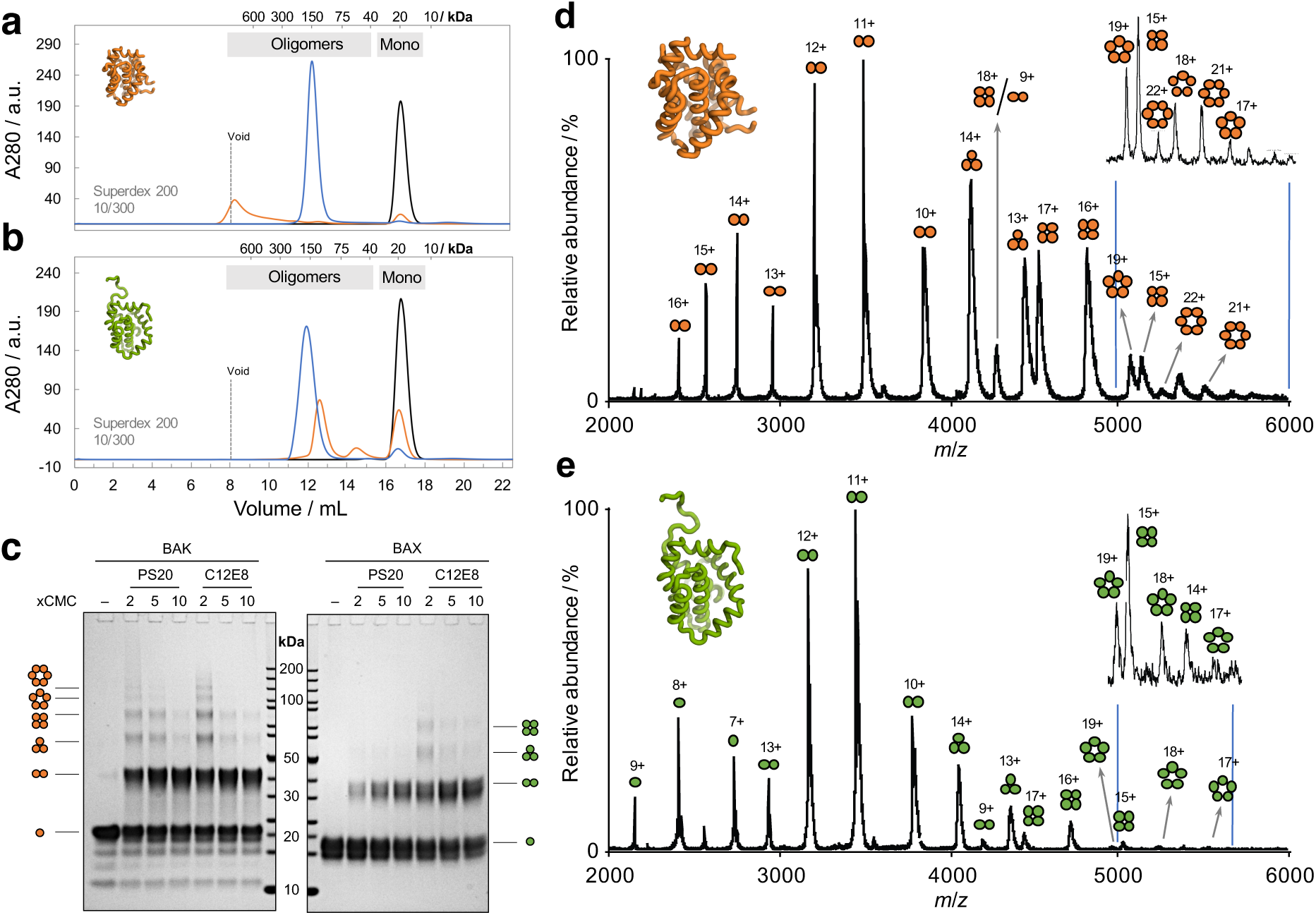
Homo-oligomerization of BAK/BAX in the presence of detergent. **a, b,** Size-exclusion chromatography of BAK (**a**) and BAX (**b**) following treatment with: no detergent (black); C12E8 at 5x critical micelle concentration (CMC, orange); PS20 at 75xCMC (blue). More detergents can be found in Extended Data Fig. 5. Some masses (based on a calibration performed with globular standards) are indicated for reference. **c,** SDS-PAGE of chemically cross-linked detergent-induced oligomers of BAK and BAX. **d, e,** Native MS of BAK (**d**) and BAX (**e**) after incubating 5 μM of monomer with PS20 (5xCMC). Insets show y-expansions for the region indicated (acquired at higher resolution). While many overlapping charge-states are evident, several unique assignments can be made (*e.g.* 6-mer 21+ and 5-mer 19+).

Use of detergents has previously been reported to result in the formation of non-physiological helix-swapped homo-dimers of BAK and BAX (Brouwer et al., 2014; Czabotar et al., 2013). However, cross-linking experiments showed that the oligomers reported here are incompatible with these structures (Extended Data Fig. 7). Moreover, we clearly observed structures larger than dimers. Importantly, Iyer et al., (2016) have shown that constraining the structure of BAK with specific disulfide staples prevented the release of cytochrome *c* from mouse embryonic fibroblast mitochondria. We found that the same disulfide mutants were incapable of oligomerization in our assay, suggesting a physiological relevance to the oligomers formed in our experiments (Extended Data Fig. 8).

We investigated the nature of these oligomers using native MS (Fig. 1d,e). Advances in the field have allowed the detection of membrane proteins in the gas-phase from detergent-solubilized complexes (Gupta et al., 2017; Laganowsky et al., 2014). Both BAK and BAX showed heterogeneous ensembles of oligomeric species, and the distributions appear to be biased towards even-numbered species. This result is reminiscent of the results obtained for BAX in supported bilayer (Subburaj et al., 2015), further reinforcing the physiological relevance of these detergent-induced oligomers. Interestingly, live-cell microscopy has revealed that BAK and BAX can form a range of pore size and shapes over the course of apoptosis (McArthur et al., 2018). Thus, the heterogeneity observed here by native MS and SEC (Extended Data Fig. 9) is probably representative of the plasticity of BAK/BAX oligomerization (Uren et al., 2017).

Having established a system capable of recapitulating oligomerization *in vitro*, we sought to address how the entire network of intermolecular interactions can lead to apoptotic pore formation, and how these interactions are regulated when all partners are present. In the ‘direct’ activation model, BH3-only proteins trigger oligomerization of BAK/BAX through transient interactions. However, we found that BH3 motifs had no appreciable affinity for either the monomeric or oligomeric states of BAK (Table 1 and Extended Data Fig. 4). Moreover, even a 10-fold excess of either BID or PUMA did not significantly affect the outcome of the oligomerization in the presence of detergent. These results challenge the ‘direct’ activation mechanism.

The presence of detergent completely altered the interaction profiles between BAK/BAX and MCL-1. Hetero-oligomers were readily recovered by SEC (Fig. 2a), contrasting with the absence of interactions observed in standard aqueous solutions. Native MS revealed the presence of hetero-dimers (BAK/BAX:MCL-1). More importantly, we found that hetero-dimer formation was at the expense of BAK/BAX self-assembly, pointing at a mechanism of competitive oligomerization (Fig. 2b,c). Thus, we infer that pore-formation can be effectively suppressed by hetero-dimerization with anti-apoptotic BCL-2 proteins. Since other anti-apoptotic BCL-2 proteins also have tight affinities for BAK_BH3_ and/or BAX_BH3,_ we anticipate this mechanism to be general (Ku et al., 2011).

**Fig. 2.**
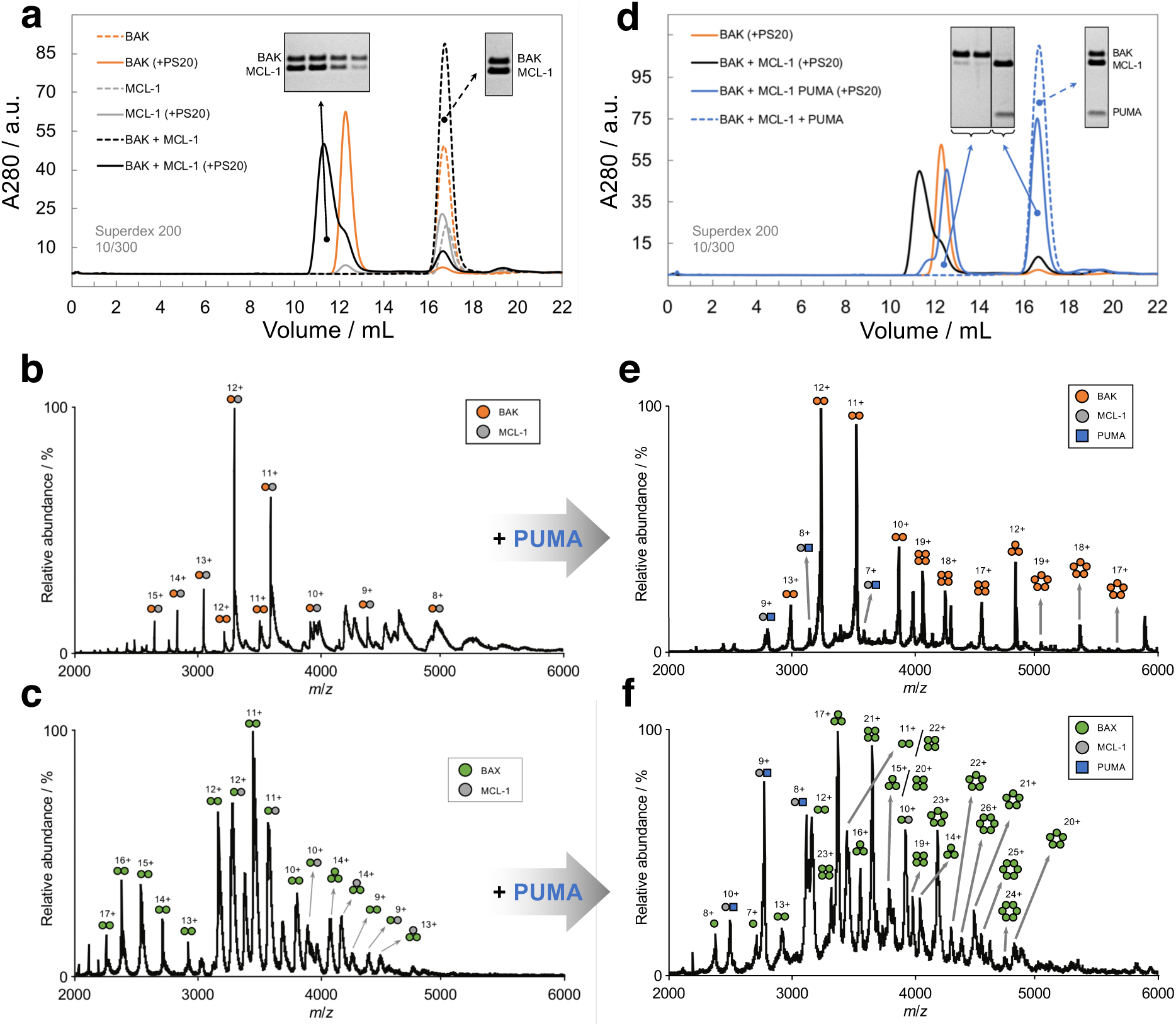
Homo-oligomerization of BAK/BAX is suppressed by hetero-dimerization with MCL-1, and recovered by addition of PUMA. **a**–**c,** Analyses of equimolar mixtures of MCL-1 with either BAK or BAX in the presence of detergent. **a,** SEC of BAK, MCL-1, and their mixture in the presence/absence of PS20. Gel insets show SDS-PAGE analyses of the corresponding peaks. **b**,**c**, Native MS of the binary mixtures of BAK (**b**) or BAX (**c**) with MCL-1 after incubation with PS20 (5xCMC). Homo-oligomerization is suppressed compared to the spectra recorded in the absence of MCL-1 (*cf.* Fig. 1d, e). **d**–**f,** Analyses of the mixtures of BAK or BAX with MCL-1 and PUMA. **d,** SEC of the ternary mixture of BAK, MCL-1 and PUMA in the presence/absence of detergent. Gel insets show SDS-PAGE analyses of the corresponding peaks. **e, f,** Native MS of the ternary mixtures of BAK (**e**) or BAX (**f**) with MCL-1 and PUMA after incubation with PS20 (5xCMC). Unlike the binary mixtures (**b, c**), formation of higher-order homo-oligomers of BAK and BAX is evident, due to PUMA scavenging MCL-1.

Suppression of pore-formation through competitive oligomerization with anti-apoptotic BCL-2 proteins is completely abrogated by BH3-only members. In the presence of PUMA, MCL-1 preferentially forms hetero-dimers with this pro-apoptotic protein, leaving BAK or BAX free to homo-oligomerize (Fig. 2d–f). These results were independent of the sequence of events; mixing the components before adding detergent, or adding MCL-1 and PUMA sequentially on pre-oligomerized BAK led to the same qualitative results. Similar results were obtained when BID was used instead of PUMA. Hence, we show unambiguously that the mechanistic role of BH3 motifs is to displace the hetero-dimers formed between BAK/BAX and MCL-1. As such, we demonstrate that their role in promoting the homo-oligomerization of BAK/BAX is entirely indirect. Together with the absence of oligomer-promoting property reported above, our results strongly support the ‘indirect’ activation model.

To gain mechanistic insights into these events, we employed kinetics (Fig. 3). The oligomerization of BAK in detergent was found to be slow, and the process captured by a single exponential decay function. Moreover, the rate of oligomerization was relatively independent of protein concentration, but was strongly affected by the quantity of detergent present (Fig. 3e,f). Thus, we postulate that the homo-oligomerization of BAK (and BAX) at the membrane is rate-limited by unimolecular processes, probably conformational re-arrangements to expose their BH3 motifs. The strong dependence of the rates on the nature and concentration of detergent also suggests that the hydrophobic environment plays a significant part in the oligomerization process (Extended Data Fig. 10). This observation might have implications for the role of membrane biophysics in the pore-formation of BAK/BAX. Interestingly, the rate of hetero-dimerization between BAK and MCL-1 in detergent was found to be very similar to the rate of homo-oligomerization under identical conditions (0.7 and 1.0 · 10^-3^ s^-1^, respectively, *cf.* Extended Data Fig. 11b). This suggests that the formations of these different assemblies are rate-limited by similar processes. Finally, the rate constant for the displacement of BAK:MCL-1 hetero-dimers by PUMA was close to the dissociation rate constant measured for MCL-1:BAK_BH3_ (Table 1 and Extended Data Fig. 11a). This result strongly suggests that the limiting step for BH3-induced oligomerization is the dissociation of BAK:MCL-1 hetero-dimers. Interestingly, the half-life measured here (∼17 min) is enticingly close to the time it takes for cytochrome *c* to be released from mitochondria in apoptotic cells (∼5 min) (Green, Goldstein, Waterhouse, Juin, & Evan, 2000). Thus, we infer that pore formation is rate-limited by the dissociation of hetero-dimers of BAK/BAX and anti-apoptotic BCL-2 proteins at the membrane.

**Fig. 3.**
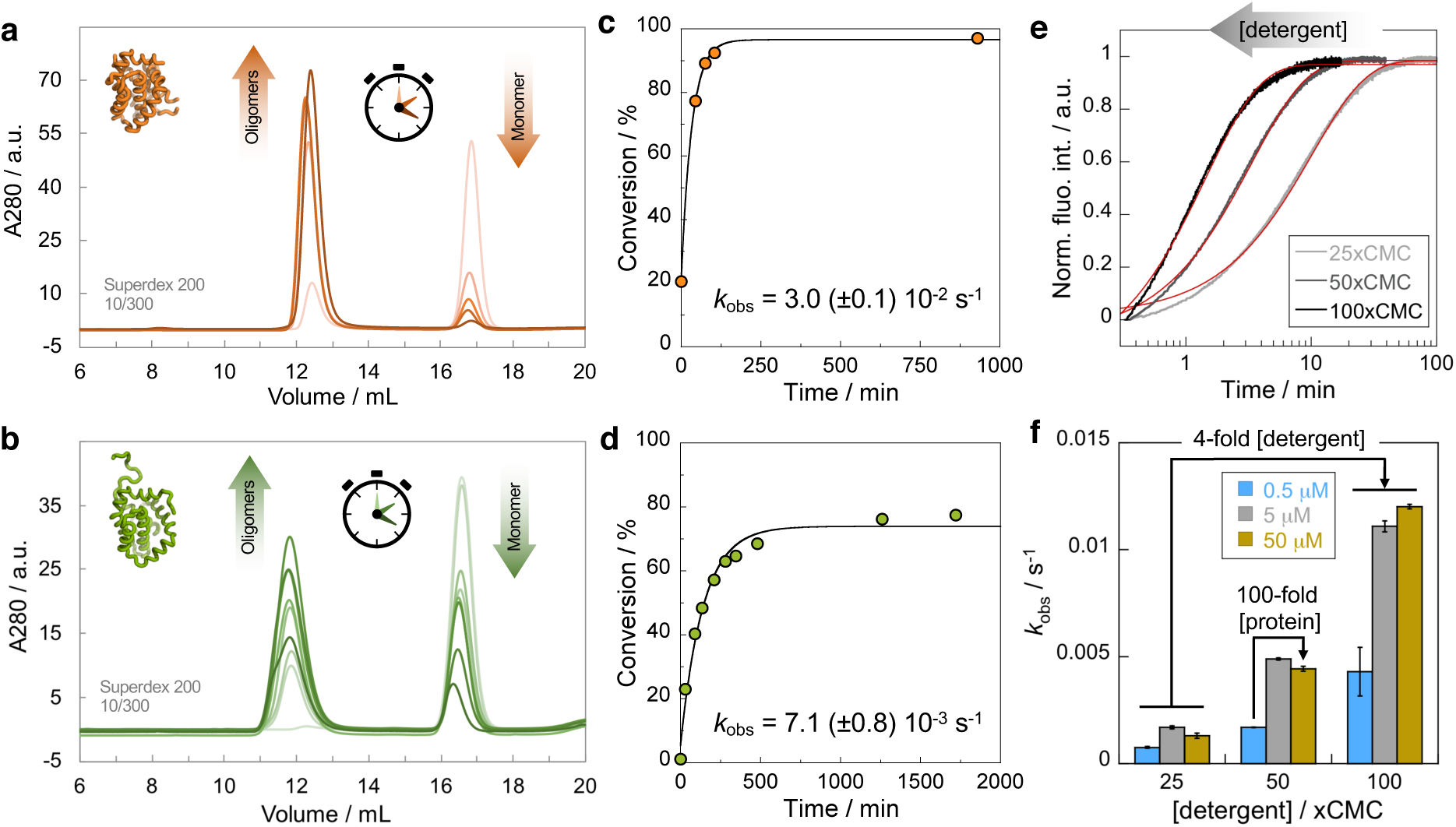
Homo-oligomerization of BAK/BAX is slow, and dominated by an apparent first-order process. Detergent-induced oligomerization of BAK (**a**) and BAX (**b**) followed by SEC time-points (18 μM monomer, 22xCMC PS20). The extents of conversation (obtained by peak integration) are plotted in (**c**) and (**d**) respectively. Solid black lines represent fits to a single exponential decay function. **e, f,** The effect of protein and detergent concentrations on the rate of BAK oligomerization was monitored spectroscopically by intrinsic tryptophan fluorescence. **e,** Example of fluorescence kinetic traces at different concentrations of PS20 and constant BAK concentration (5 μM). Solid red lines are fits to single exponential decay functions. **f**, Detergent concentration has a much greater effect on the rate of oligomerization than protein concentration. BAK oligomerization was monitored by fluorescence over 4-fold and 100-fold ranges of PS0 and protein concentrations respectively. Error bars represent standard deviations (n = 2).

**Fig. 4.**
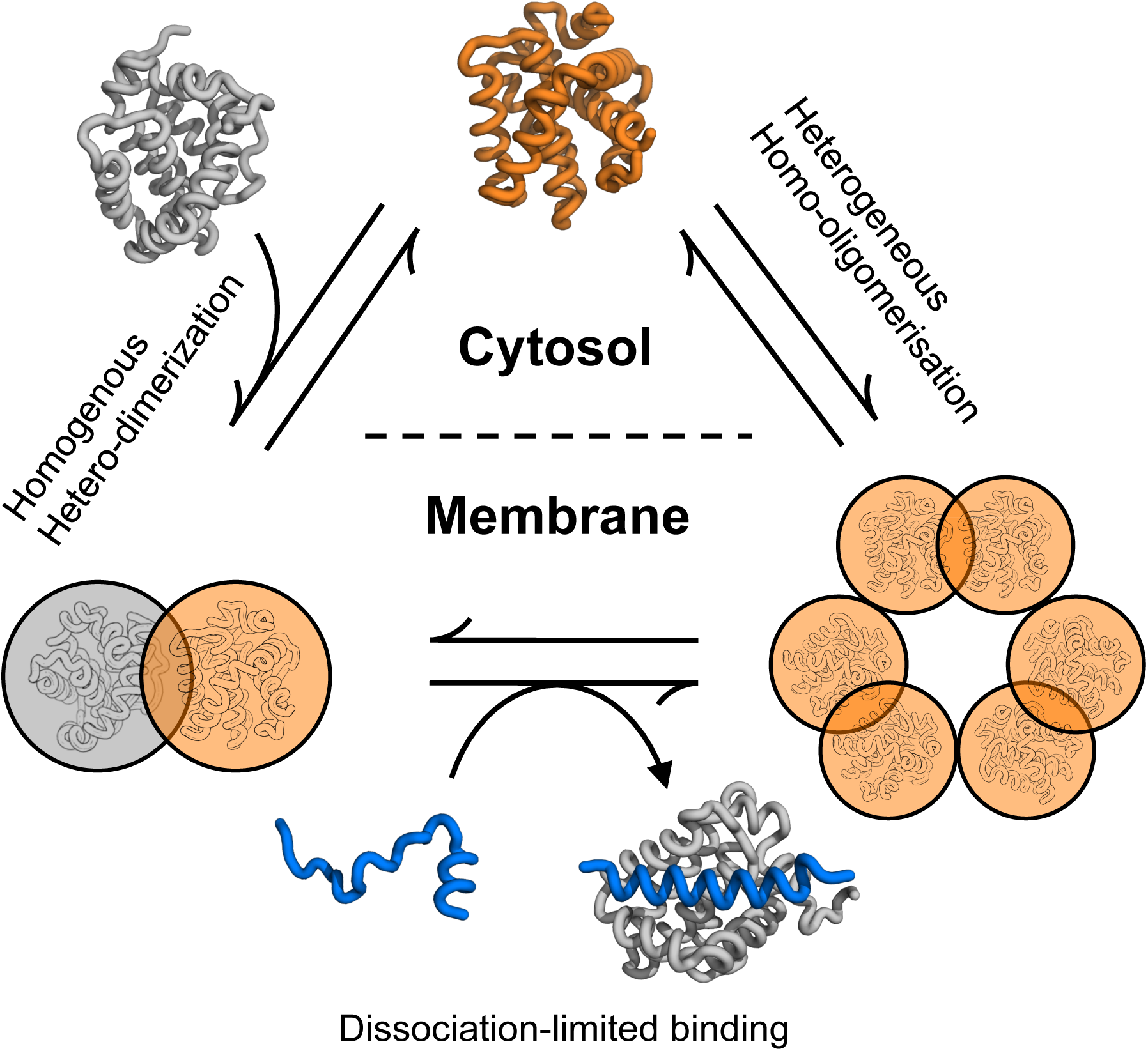
Putative schematic of competitive oligomerization occurring at the membrane. BAK (and BAX) are monomeric in the cytosol, but spontaneously homo-oligomerize at the membrane. These structures are heterogeneous (possibly based on dimer units). The presence of MCL-1 at the membrane ensures specific hetero-dimerization at the expense of homo-oligomerization. The presence of PUMA passively disrupts BAK:MCL-1 hetero-dimers (rate-limited by the dissociation of the complex), leaving BAK (and BAX) free to oligomerize. BAK, MCL-1 and PUMA are shown in orange, grey, and blue respectively. Structures based on PDB 2MHS (MCL-1), 2YV6 (BAK), and 2ROC (PUMA:MCL-1).

Our results clarify a long-standing debate regarding the physical interactions and mechanistic details underlying BCL-2 regulation. Our findings demonstrate that a membrane-like hydrophobic environment is sufficient to trigger spontaneous homo-oligomerization of BAK/BAX, and that BH3 motifs do not participate in this process. Thus, the membrane appears to be the true ‘activator’ of oligomerization. This conclusion is supported by a recent cellular study which reported that knocking-out all BCL-2 proteins except BAK and BAX resulted in spontaneous apoptosis (O’neill, Huang, Zhang, Chen, & Luo, 2016). We further demonstrate that the role of anti-apoptotic BCL-2 proteins is to suppress the homo-oligomerization of BAK/BAX by forming hetero-dimers with them, and BH3-only proteins exercise their pro-apoptotic activity only indirectly—by displacing BAK/BAX from their anti-apoptotic ‘chaperones’, and leaving them free to oligomerize. The work presented here lays the foundation for future quantitative biophysical characterizations of the apoptotic interactome, with implications for the understanding of pathological conditions and new treatment paradigms.

## Acknowledgments

This work was supported by the Wellcome Trust (WT095195). J.C. is a Wellcome Trust Senior Research Fellow. B.I.M.W. was supported by a scholarship from the Cambridge Trust. K.G. and C.V.R. are grateful for support of the European Research Council grant no. 69551-ENABLE and a Wellcome Trust Investigator Award (104633/Z/14/Z).

## Author Contributions

B.I.M.W. and T.O.C.K. produced the proteins. B.I.M.W. performed and analyzed all biophysical experiments. K.G. and B.I.M.W. performed and analyzed native MS. B.I.M.W., K.G., C.V.R., and J.C. conceived and planned the investigations and wrote the manuscript.

## Author Information

The authors declare no competing financial interests. Correspondence and requests for materials should be addressed to J.C. or C.V.R.

## Data Availability

All data supporting the findings of this study are available from the corresponding authors upon request.

## Methods

### Constructs

Coding DNA sequences for human BAK, BAX and MCL-1 were ordered from Genscript (codon-optimized for *Escherichia coli*), and sub-cloned into expression vectors by scarless insertion using the In-Fusion kit (Takara). Cysteine residues were mutated to serines to avoid the use of reducing agents. MCL-1 (UniProt:Q07820, residues 168–327, C286S) was sub-cloned into a modified version of the pRSET A vector containing a N-terminal hexahistidine-tag followed by a thrombin cleavage site. The disordered N-terminus, and the C-terminal transmembrane regions were excluded, in line with literature reports. An extra GS remained at the N-terminus following proteolytic tag removal. Constructs for BAK (UniProt:Q16611, residues 16–185, C166S) and BAX (UniProt:Q07812, residues 1–171, C62S, C126S) were sub-cloned into the pTXB1 vector (New England Biolabs), which contains a C-terminal intein, followed by a chitin binding domain. BAK and BAX constructs were designed to match literature reports of these proteins. In both cases, the C-terminal transmembrane helix was removed to aid solubility. In the case of BAK, a short disordered segment at the N-terminus was also excluded. Cysteine mutants of BAK were obtained by site-directed mutagenesis of the parental construct. All cloning results were confirmed by sequencing.

### Protein expression and purification

MCL-1: Plasmids were transformed into C41(DE3) cells, and grown overnight at 37 °C on 2xTY-agar plates containing ampicillin (100 μg/mL). Pre-cultures (5–10 mL) were prepared from scrapes of these plates, and used to inoculate 1 L cultures (LB). Cells were grown at 37 °C until they reached an OD_600_ of about 0.6. Protein expression was induced by addition of IPTG (1 mM final concentration), and expression was carried out overnight at 18 °C. Cells were harvested by centrifugation, the pellet re-suspended in PBS buffer (10 mM sodium phosphate, 137 mM NaCl, 3 mM KCl, pH 7.4) containing 25 mM imidazole, and sonicated on ice. Debris were cleared by centrifugation, and the proteins purified from the supernatant by binding to Ni-NTA resin. The protein was released by addition of PBS containing 500 mM imidazole and buffer-exchanged into 20 mM Tris, 150 mM NaCl, 5 mM CaCl_2_, pH 7.5. Cleavage of the hexahistidine-tag was performed overnight at room temperature using thrombin from bovine serum (Sigma). MCL-1 was further purified by ion-exchange chromatography on a HiTrap SP HP cation-exchange column (GE Healthcare) using a 10 mM HEPES pH 7.5 ± 1 M NaCl buffer system (stepped gradient; 0–9% over 20 mL, 9–13% over 25 mL, 13–20% over 20 mL). Clean fractions were pooled, and further purified by SEC on a Superdex 75 26/600 (GE Healthcare) equilibrated in 50 mM sodium phosphate pH 7.0.

BAK and BAX: Plasmids were transformed into BL21(DE3) (BAK) or C41(DE3) (BAX) cells, and grown overnight at 37 °C on 2xTY-agar plates containing ampicillin (100 μg/mL). Pre-cultures made from these plates were used to inoculate 1 L cultures (LB). Cells were grown at 37 °C until they reached an OD_600_ of about 0.6. Protein expression was induced by addition of IPTG. BAK expression was carried out at 37 °C for 4 h following induction with 1 mM IPTG. BAX was induced with 0.1 mM IPTG, and the expression performed overnight at 28 °C. Cells were harvested by centrifugation, the pellet re-suspended in HEPES buffer (20 mM HEPES, 100 mM NaCl, 1 mM EDTA, pH 7), and sonicated on ice. Debris were cleared by centrifugation, and the proteins were purified from the soluble fraction by binding to chitin resin (New England Biolabs) at 4 °C. Removal of the tag was achieved by self-cleavage of the intein, which was induced by the addition of 50 mM DTT overnight at room temperature. Cleaved proteins were eluted from the column, concentrated, and further purified by SEC on a Superdex 75 26/600 equilibrated in 50 mM sodium phosphate pH 7.0.

PUMA_BH3_ used for out-competition experiments was produced recombinantly as a GB1 fusion. The synthetic gene was ordered from Genscript and sub-cloned into pRSET A. The construct contained a N-terminal hexahistidine-tag followed by GB1 fused to PUMA_BH3_ (UniProt:Q9BXH1, residues 125–161, M144A) via a thrombin cleavage site. Expression and purification protocols were broadly similar to MCL-1. In brief, expression was carried out at 37 °C for 4 h in C41(DE3) or C41(DE3)pLysS cells. After sonication, the construct was purified from the soluble fraction by nickel affinity chromatography, and the peptides was cleaved from its His_6_-GB1 tag overnight at room temperature using thrombin. The released peptide was first purified by anion-exchange chromatography (HiTrap Q HP, GE Healthcare) with a 20 mM Tris pH 8.0 ± 1 M NaCl buffer system (0–20% over 50 mL, elution at about 15%), followed by SEC on a Superdex 30 26/600 (GE Healthcare) equilibrated in 50 mM sodium phosphate pH 7.0.

Protein purities were assessed by SDS-PAGE and coomassie staining, and identities were confirmed by LC-MS (MCL-1: 18409.52(±4.33), theoretical 18409.74 Da; BAK: 19213.01(±8.55), theoretical 19217.64 Da; PUMA_BH3_: 4441.93(±0.50) Da, theoretical 4442.75 Da). The mass of BAX was confirmed by native MS (within 1 Da of its theoretical value of 18933.65 Da).

Dye-labelled peptides of the BH3 motifs were ordered from Biomatik as 35 amino acid long peptides composed of the BH3 sequence (15 residues), plus 10 flanking residues on both sides. Each construct also contained a N-terminal 5-carboxytetramethylrhodamine (TAMRA) dye for extrinsic fluorescence experiments. All sequences were based on gene products from *Homo sapiens*: PUMA_BH3_ (UniProt: Q9BXH1, residues 127–161, M144A); BID_BH3_ (UniProt:P55957, residues 76–110); BAK_BH3_ (UniProt:Q16611, residues 64–98, P64A); BAX_BH3_ (UniProt:Q07812, residues 49–83, P49G and C62A); MCL-1_BH3_ (UniProt:Q07820, residues 199–233).

### Detergent-induced oligomerization

Stock solutions of detergents were prepared in 50 mM sodium phosphate pH 7.0 or 100 mM ammonium acetate, pH 7.0 (for native MS). PS20 (polyoxyethylene (20) sorbitan monolaurate, Fisher Scientific) stock solutions were prepared at a concentration of 80 mM (9% v/v, 1000xCMC) by adding 90 μL of neat liquid to 910 μL of buffer. C12E8 (octaoxyethylene monododecyl ether, Anatrace) was purchased as a 25% w/w stock in water (∼470 mM). Working stock solutions were prepared at 90 mM (1000xCMC) by adding 192 μL of stock to 808 μL of buffer. C8E4 (tetraoxyethylene monooctyl ether, Anatrace) was purchased as a 50% w/w stock in water (∼1.6 M). Working stock solutions were prepared at 80 mM (10xCMC) in buffer, or used neat. OGP (octyl b-D-glucopyranoside, Sigma-Aldrich) stock solutions were prepared at a concentration of 100 mM (4xCMC) by dissolving 58.5 mg in 2 mL of buffer. The oligomerization of BAK and BAX were induced by adding detergents from stock solutions to the final concentrations indicated in the text (expressed as multiples of their CMC’s). Typical protein concentrations for these experiments were 10-20 μM, and samples were incubated from a few hours to overnight to ensure that the systems reached equilibrium (incubation times were adjusted depending on the nature of the detergent, and its concentration).

### Chemical cross-linking

Detergent-treated BAK/BAX oligomers were cross-linked with a mixtures of EDC (1-ethyl-3-(3-dimethylaminopropyl)carbodiimide hydrochloride, Thermo Scientific) and BS3 (bis(sulfosuccinimidyl)suberate, Thermo Scientific) at a molar ratio of 1:300:300 (protein:EDC:BS3). Cross-linker stock solutions were prepared in 50 mM sodium phosphate pH 7.0, and used directly. Proteins (20 μM of monomer) were pre-incubated in detergent, followed by cross-linking at room temperature for 2 h. Results were analyzed by SDS-PAGE and coomassie straining.

### Size-exclusion chromatography

Analytical size-exclusion chromatography experiments were performed on a Superdex 200 Increase 10/300 GL column (GE Healthcare) equilibrated in 50 mM sodium phosphate pH 7.0. Injection volumes were between 100–500 μL and elutions were performed at 0.75 mL/min. Chromatograms were recorded by measuring the absorbance at 280 nm. Proteins were pre-incubated in detergent prior to injection. Typical protein concentrations were 10–20 μM, and detergent concentrations are indicated in the text as multiple of their CMC’s. The calibration of the column was performed using globular chromatographic standards (GE Healthcare), and found to be: log(MW/Da) = – 0.194 x V_elution_ + 7.57.

### Disulfide cross-linking

Oxidation of BAK cysteine double mutants was performed using the redox catalyst Cu^II^(1,10-phenanthroline)_3_ (Kobashi, 1968). Stock solutions (25 mM, prepared from copper sulfate and 1,10-phenantroline in a 4:1 water:ethanol mixture) were added to protein solutions at a final concentration of 0.5 mM, followed by incubation on ice for 30–45 min. The reactions were quenched by addition of EDTA (2 mM final concentration), followed by overnight dialysis at room temperature. Results were analyzed by SEC or non-reducing SDS-PAGE. Control experiments to assess the effect of cysteine mutations on oligomerization were obtained by reducing the respective disulfide mutant with 50 mM DTT for 10 min at room temperature prior to SEC analysis. The oxidation step (disulfide stapling) was performed either before, or after, the addition of detergent (as detailed in the text).

### Spectroscopic analysis of the oligomers

Spectroscopic signatures associated with the monomeric and detergent-treated oligomeric states of BAK and BAX were analysed by far-UV CD spectroscopy and intrinsic tryptophan (and tyrosine) fluorescence at protein concentrations between 5 and 10 μM. CD spectra were recorded on a Chirascan instrument (Applied Photophysics) between 200 and 250 nm (1 nm intervals) and adaptive sampling. Fluorescence emission spectra were recorded on a Cary Eclipse Fluorescence Spectrophotometer (Agilent) by exciting at 280 nm and recording the fluorescence intensity between 300 and 400 nm (1 nm increments). All spectra were buffer-subtracted.

### Oligomerization kinetics

Oligomerization kinetics in the presence of detergents was assessed by intrinsic tryptophan fluorescence. Reactions were monitored on a Cary Eclipse Fluorescence Spectrophotometer (Agilent) by exciting at 280 nm, and measuring changes in fluorescence intensity at 330 nm. Reactions were performed in 50 mM sodium phosphate pH 7.0, and the oligomerization initiated by addition of detergent from 10x stocks using a pipette. The dead time between the addition of detergent and the start of the data acquisition were accounted for by modifying the timebase during analysis. Measurements were performed in duplicates, traces individually fitted to a single exponential decay function, and the observed rates averaged :

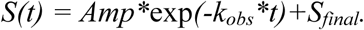

### Association kinetics

Association rate constants for BH3 peptides and either MCL-1 or BAK were obtained using stopped-flow kinetic measurements under pseudo-first order conditions. Experiments were recorded on either a SX18 or SX20 stopped-flow spectrophotometer (Applied Photophysics). Reactions were monitored by either following the change in intrinsic fluorescence (exciting at 280 nm, and using a 320 nm longpass filter), or extrinsic fluorescence (TAMRA, exciting at 555 nm, and using a 570 nm longpass filter). In some cases, the signal change observed from fluorescence intensity was poor, and fluorescence anisotropy was recorded instead (using a FP1 fluorescence polarisation accessory, Applied Photophysics). Typical concentrations of TAMRA-labeled peptides were 100–500 nM (final), and the partner protein was present in at least 10-fold excess. For each condition (pair of concentrations), multiple association traces were recorded (10-40), the data averaged, and the result fitted to a single exponential decay function. Some association kinetics between MCL-1 and BH3 motifs were performed using un-labeled peptides. These reactions were performed under reverse-pseudo-first order conditions (peptide in excess), and also fitted to a single exponential decay function. The gradient of the line between *k*_obs_ and the concentration of excess partner gave *k*_on_

### Dissociation kinetics

Dissociation rate constants for complexes between BH3 motifs and MCL-1 were obtained by performing out-competition experiments. Complexes of MCL-1 and TAMRA-labeled BH3 peptides were pre-formed by mixing equimolar amounts of the components (typical concentrations were 5 μM). A solution of the out-competitor (PUMA_BH3_, un-labeled) was placed in a fluorescence cuvette, and irreversible dissociation was initiated by diluting the complex into the un-labeled BH3 peptide solution. Data were collected by exciting TAMRA at 555 nm and measuring the change in fluorescence intensity at 575 nm. In certain cases, loss of fluorescence polarisation (obtained using a manual fluorescence polarisation accessory set to V/V) gave a better signal-to-noise ratio, and was used instead of fluorescence intensity. Data were fitted to a single exponential decay function. For long traces, a linear drift term was added to account for photobleaching of the dye. Experiments were repeated in the presence of different excess quantities of out-competitor (50–400 molar excess over complex, two different excess concentrations for each pair). Irreversible dissociation was confirmed by the absence of a dependence of *k*_obs_ on the fold-excess of competitor. Concentration-independent values were averaged, giving *k*_off_. For some out-competition dissociation reactions, the rates were too fast to be measured by manual mixing on a fluorescence spectrophotometer, in which cases stopped-flow kinetic measurements using 1:10 volume mixing were performed. General principles were identical to the experiments performed by manual mixing, but the reduction in dead time allowed fast dissociation events to be observed.

### Equilibrium binding

Equilibrium binding dissociation constants between BAK and BH3 motifs were obtained from binding isotherms by monitoring the fluorescence anisotropy of TAMRA-labeled peptides as a function of BAK concentration. Experiments were performed on a Cary Eclipse Fluorescence Spectrophotometer (Agilent) using a manual polarization accessory, exciting the dye 555 nm and recording the fluorescence at 575 nm. Binding curves were prepared by serial dilutions of BAK in the presence of a constant concentration of dye-labeled peptide (1 μM). BAK was concentrated using centrifugal concentrators, and its concentration was determined spectroscopically on a Cary 60 UV-Vis spectrophotometer (Agilent). Binding curves were prepared by serial dilutions of BAK (1:1 volumes). For each point, the fluorescence intensity in each polarization plane was measured in triplicate, and averaged. These readings were converted to anisotropy values:

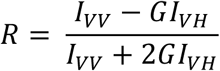

where *R* is the fluorescence anisotropy, *I*_VV_ and *I*_VH_ are the vertically- and horizontally-polarized components of the fluorescence intensity, and *G* = *I*_HV_/*I*_HH_ is a correction factor that accounts for the instrument’s differential detection sensitivities in the vertical and horizontal polarization planes (where *I*_HV_ and *I*_HH_ have analogous meanings to *I*_VV_ and *I*_VH_). To account for changes in fluorescence intensity upon binding, the anisotropy was corrected using the following expression:

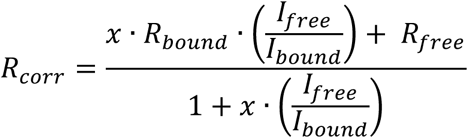

where *R*_corr_ represents the corrected anisotropy, *R*_free_ and *R*_bound_ the anisotropy of free and bound peptide respectively, *I*_free_ and *I*_bound_ the total fluorescence intensities of the free and bound states respectively (*I* = *I*_VV_ + 2*I*_VH_), and *x* = (*R* – *R*_free_)/(*R*_bound_ – *R*).

Equilibrium dissociation constants (*K*_d_) were obtained by fitting the corrected anisotropy signal as a function of BAK concentration to a 2-state hetero-dimerisation model:

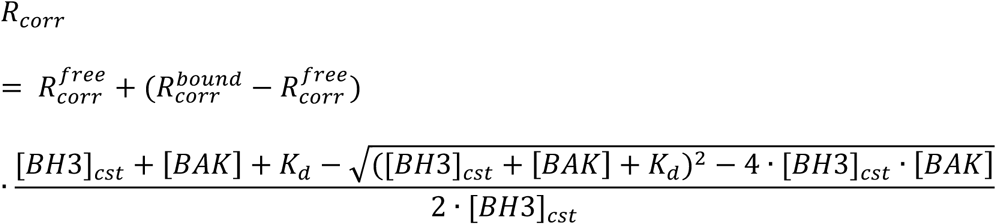

where 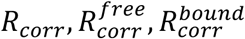 represent the observed corrected anisotropy, the corrected anisotropy of the free peptide and the corrected anisotropy of the bound peptide respectively. [BH3]_cst_ represent the concentration of labeled peptide (constant), [BAK] the concentration of BAK (the independent variable) and *K*_d_ the equilibrium dissociation constant.

### Native mass spectrometry

Protein stocks were buffer-exchanged into 100 mM ammonium acetate pH 7.0 by SEC on a Superdex 75 10/300. Mixtures of proteins were assembled in the buffer to final concentrations of 5 μM (total, equimolar distributions), before adding PS20 (5xCMC). Oligomerization was left to proceed at room temperature for at least 3 h before analyzing the samples. All spectra were acquired on a modified Q Exactive hybrid quadrupole-Orbitrap mass spectrometer (Thermo Scientific)(Fort et al., 2018) coupled with an offline source, at a HCD cell pressure of 7 mL/min of argon, and an HCD energy of 40 eV. The data were analyzed with Xcalibur 4.1 (Thermo Scientific).

## Extended Data Figures

**Extended Data Fig. 1.**
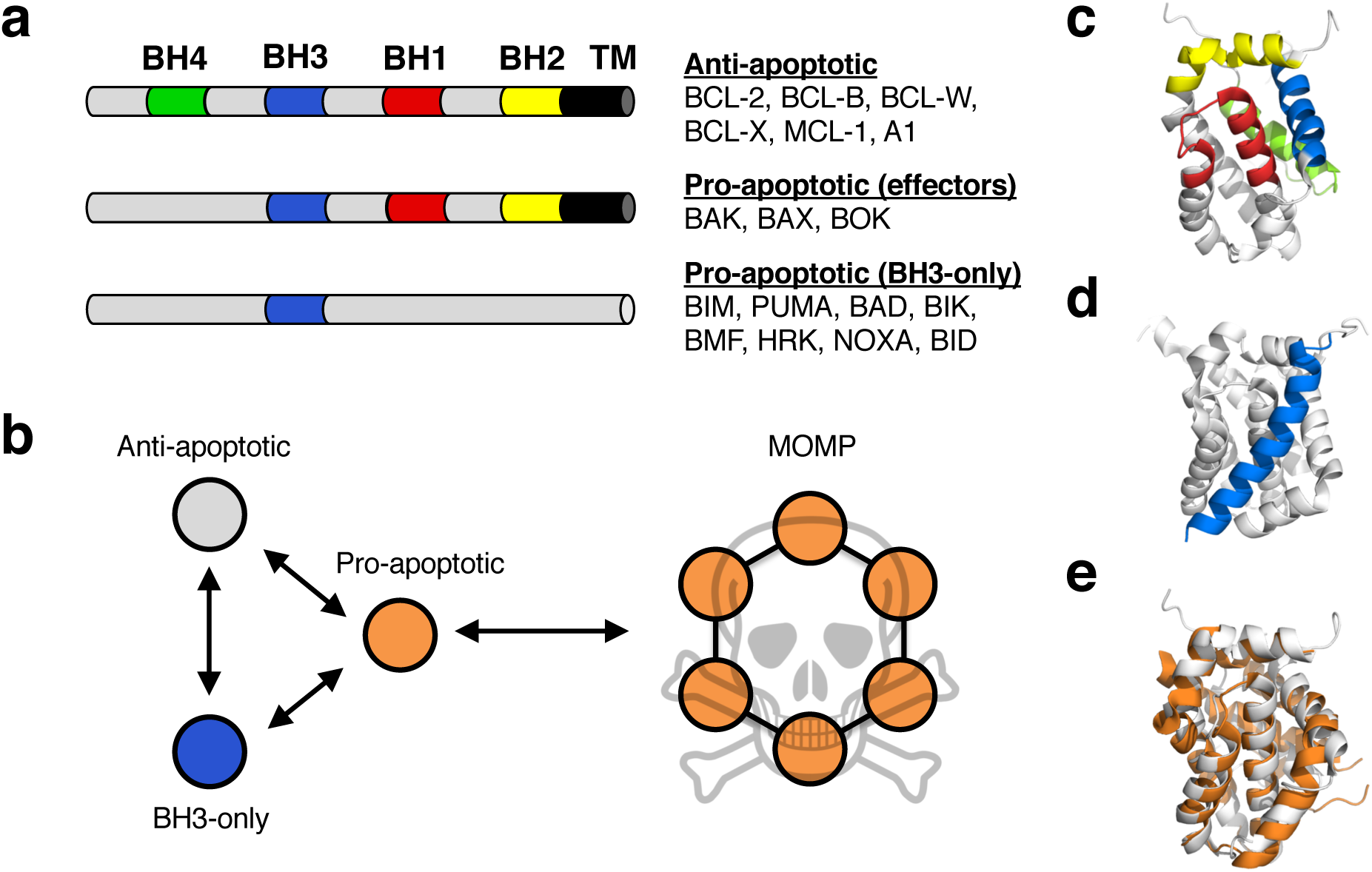
The BCL-2 family. **a,** Sequence and functional classification of BCL-2 proteins. Most multi-motifs BCL-2 have a putative transmembrane (TM) helix. **b,** General putative interaction network controlling the oligomerization of BAK/BAX leading to mitochondrial outer-membrane permeabilization (MOMP). **c,** Structure of BCL-2 (PDB:1G5M) with its BH motifs highlighted (color scheme identical to (**a**)). **d,** Structure of MCL-1 bound to PUMA_BH3_ at the canonical groove (PDB:2ROC). **e,** Structural alignment of BCL-2 (PDB:1G5M) and BAK (PDB:2YV6) showing the structural similarities between pro- and anti-apoptotic BCL-2 proteins. The orientations of the structures in **c**–**e** are identical.

**Extended Data Fig. 2.**
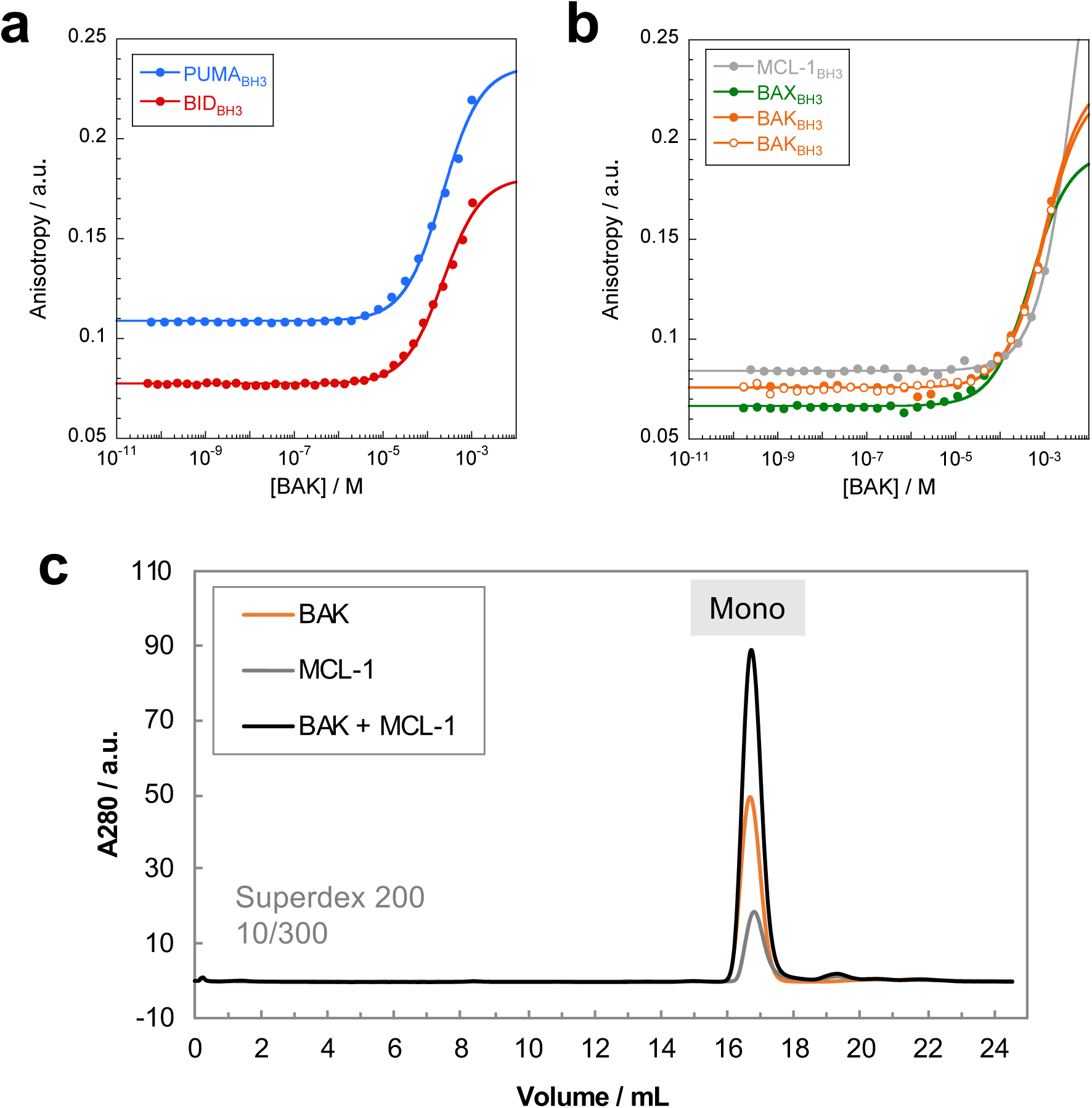
Interactions between BAK, BH3 peptides and MCL-1 in the absence of detergent. **a, b,** Binding isotherms for BAK and TAMRA-labeled BH3 peptides from: **a,** BH3-only proteins and **b,** multi-motifs BCL-2 proteins. Binding was assessed by fluorescence anisotropy of the dye, and solid lines represent fits to a 2-state binding model (*K*_d_ values are reported in Table 1). In (**b**) the results from two independent experiments are shown for BAK_BH3_. **c,** Size-exclusion analysis of a mixture of BAK and MCL-1 (in the absence of detergent) shows no oligomerization after overnight incubation at 25 °C. Protein concentrations were 10 μM each.

**Extended Data Fig. 3.**
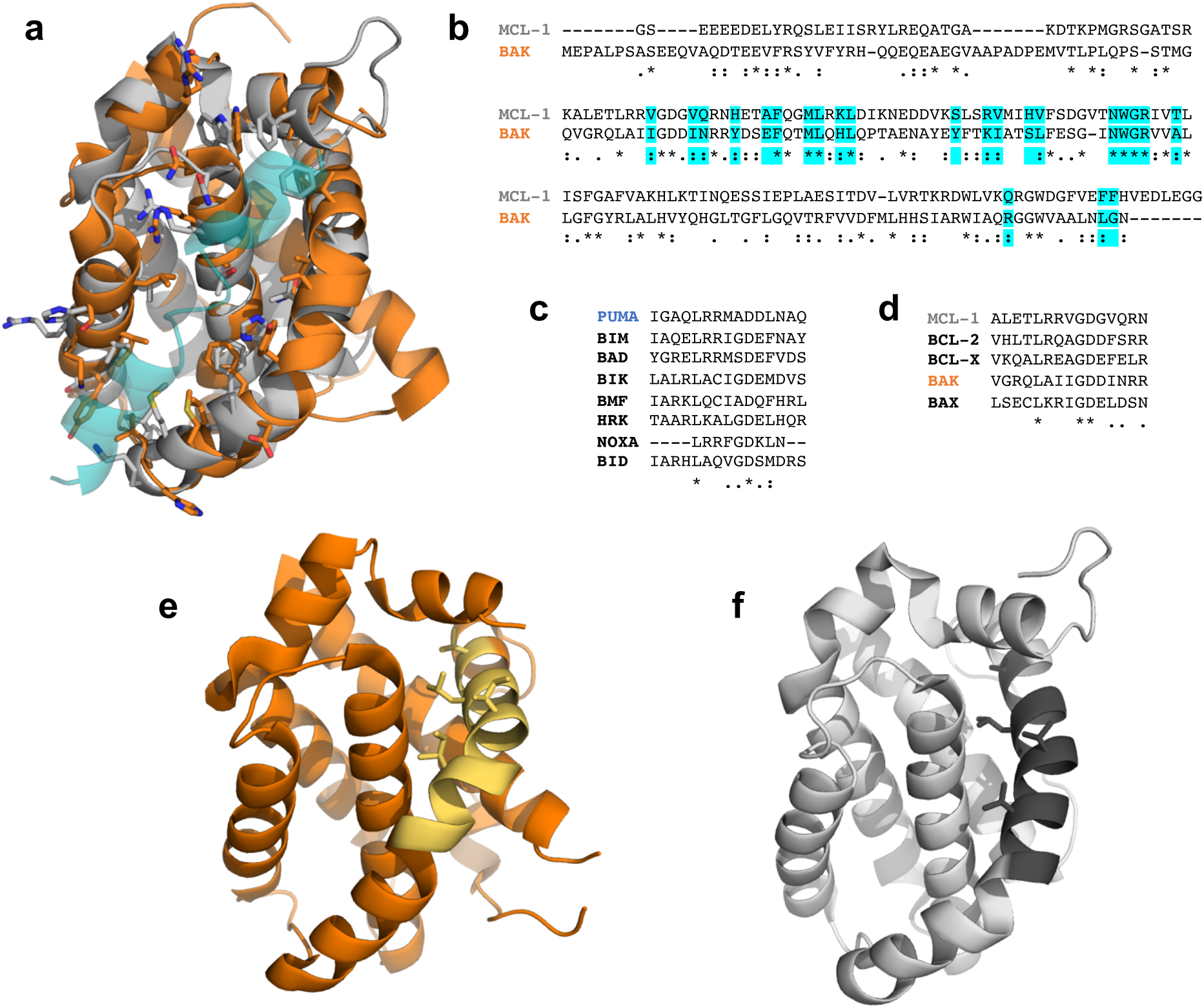
Homology between BAK and MCL-1. **a,** Structural comparison between BAK (orange, PDB:2M5B) and MCL-1 (grey, PDB:2MHS). Interface residues lining the BH3 groove are highlighted in sticks representation. The stapled BID (bound to BAK) is shown in transparency to aid visualization of the groove. The RMSD over 131 Cα is 2.76 Å. **b,** Sequence alignment of BAK and MCL-1. Residues at the interface of the BH3 groove are highlighted in cyan. The identities over the full sequence and the interface are 19.9% and 34.8% respectively. **c,** BH3 motifs from BH3-only proteins. These are part of larger, intrinsically disordered proteins. **d,** BH3 motifs from anti-apoptotic (MCL-1, BCL-2, BCL-X) and pore-forming (BAK and BAX) BCL-2 proteins. These sequences are part of folded regions within their respective proteins. **e, f,** BH3 regions of BAK (**e**, PDB:2YV6) and MCL-1 (**f**, PDB:2MHS) are highlighted as a different shade within their respective structures. Residues composing the hydrophobic side of the amphipatic BH3 helix (sticks representation) face inwards, and are inaccessible for interactions in these monomeric folded states.

**Extended Data Fig. 4.**
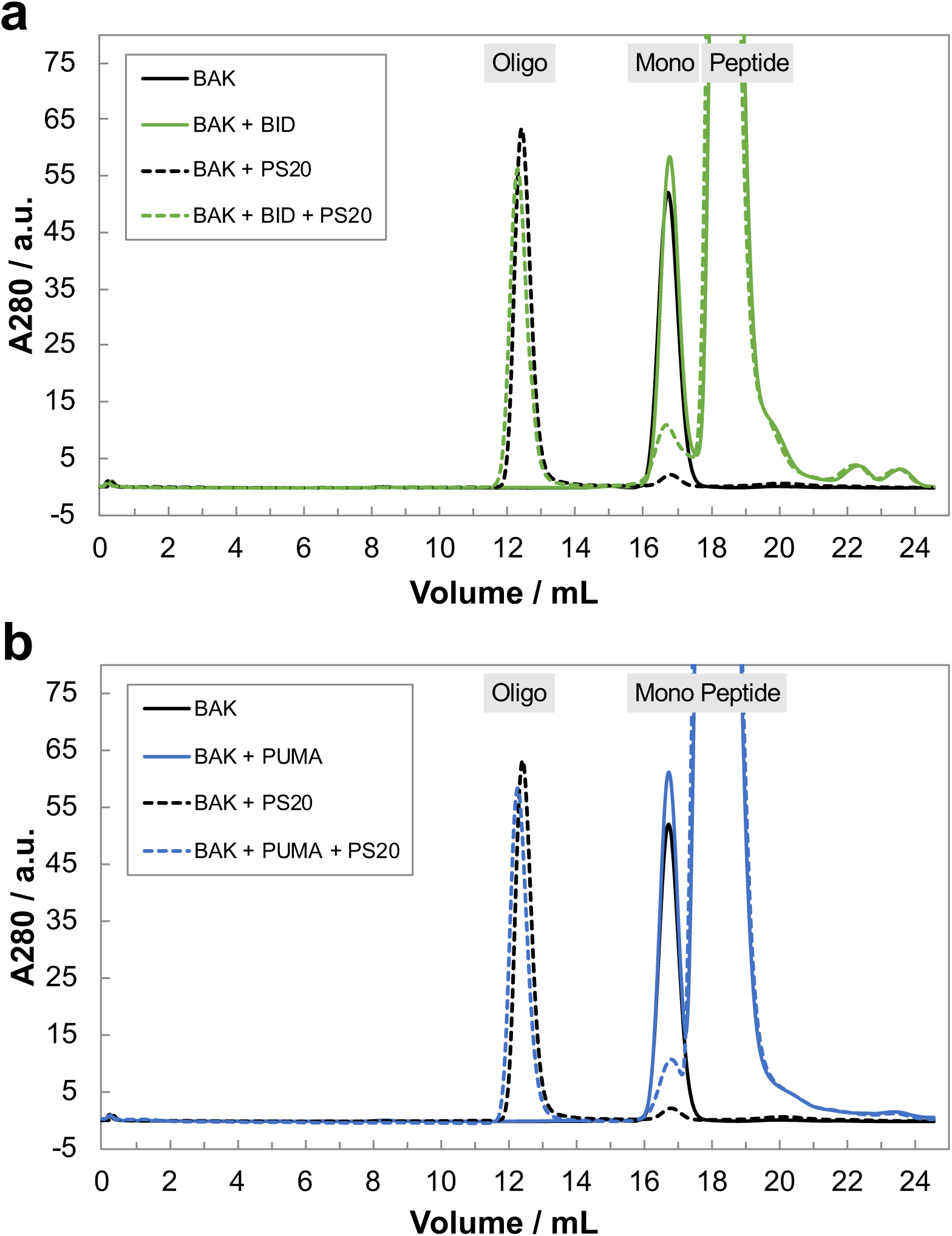
BH3 peptides do not promote BAK oligomerization. Effect of (**a**) BID_BH3_ and (**b**) PUMA_BH3_ on BAK after two days of incubation at 25 °C. These BH3 motifs are spectators, indicated by a lack of binding, and an unaffected oligomerization profile. Concentrations of protein and peptides were 10 μM and 100 μM respectively. Experiments were performed in the presence/absence of PS20 (20·CMC). Absence of peptide in the oligomeric peak was confirmed by a lack of absorbance at 555 nm (the absorbance maximum of the TAMRA dye).

**Extended Data Fig. 5.**
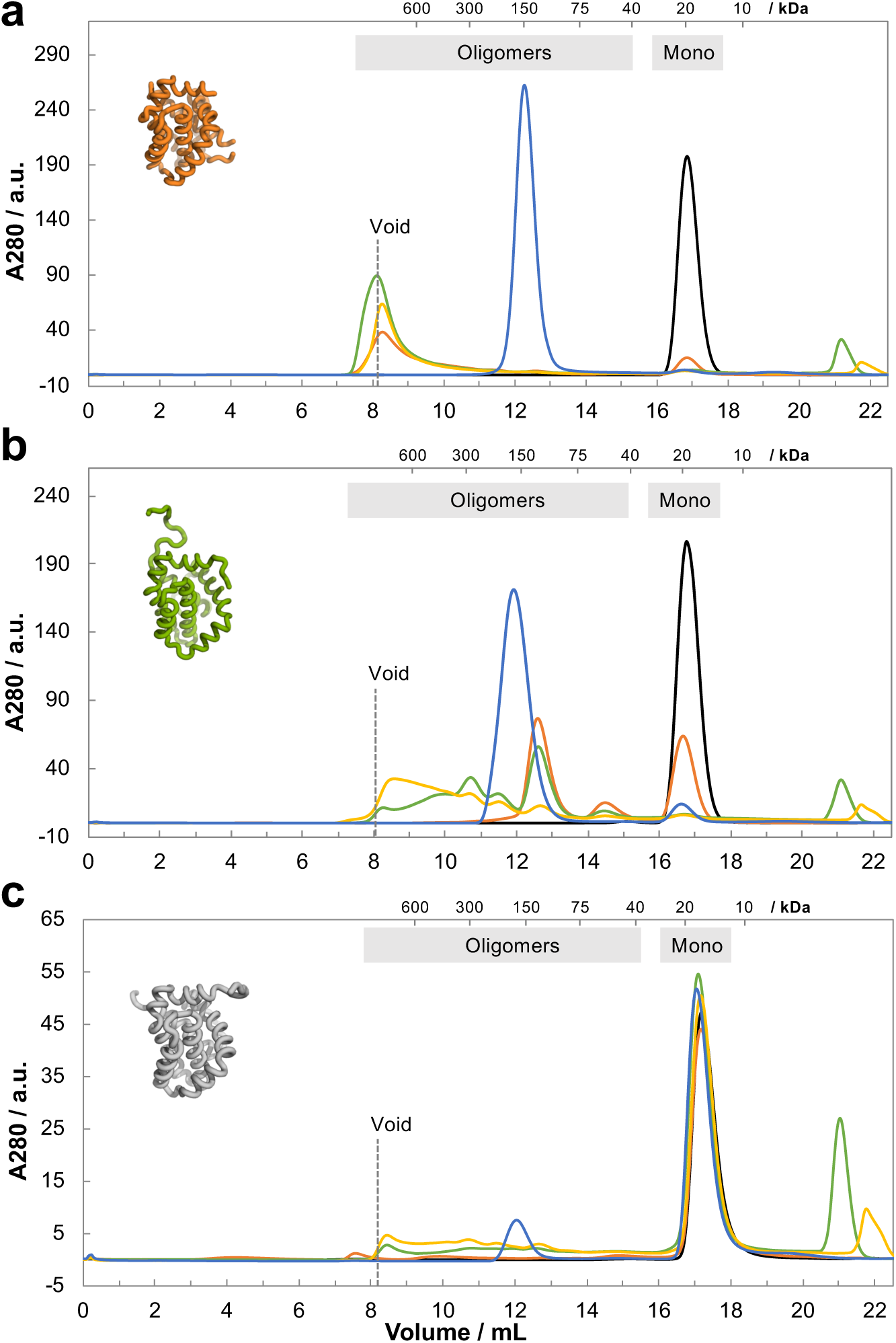
Effect of detergents on BAK/BAX and anti-apoptotic MCL-1. **a,** BAK. **b,** BAX. **c,** MCL-1. Protein concentrations were 50 μM for BAK/BAX, and 20 μM for MCL-1. Conditions were: no detergent (black); C12E8 at 5·CMC (orange); C8E4 at 2·CMC (green); OGP at 2·CMC (yellow); PS20 at 75·CMC (blue). The peak at ∼12 mL in (**c**) is solely due to the absorbance of the detergent micelle, and does not contain any protein (confirmed by SDS-PAGE). The smaller detergents OGP and C8E4 appeared to partially unfold the proteins (especially MCL-1); indicated by peaks with anomalous late elution volumes (after the monomers). Some masses (determined by calibration with globular standards) are reported above each plot for reference. All experiments were performed on a Superdex 200 10/300.

**Extended Data Fig. 6.**
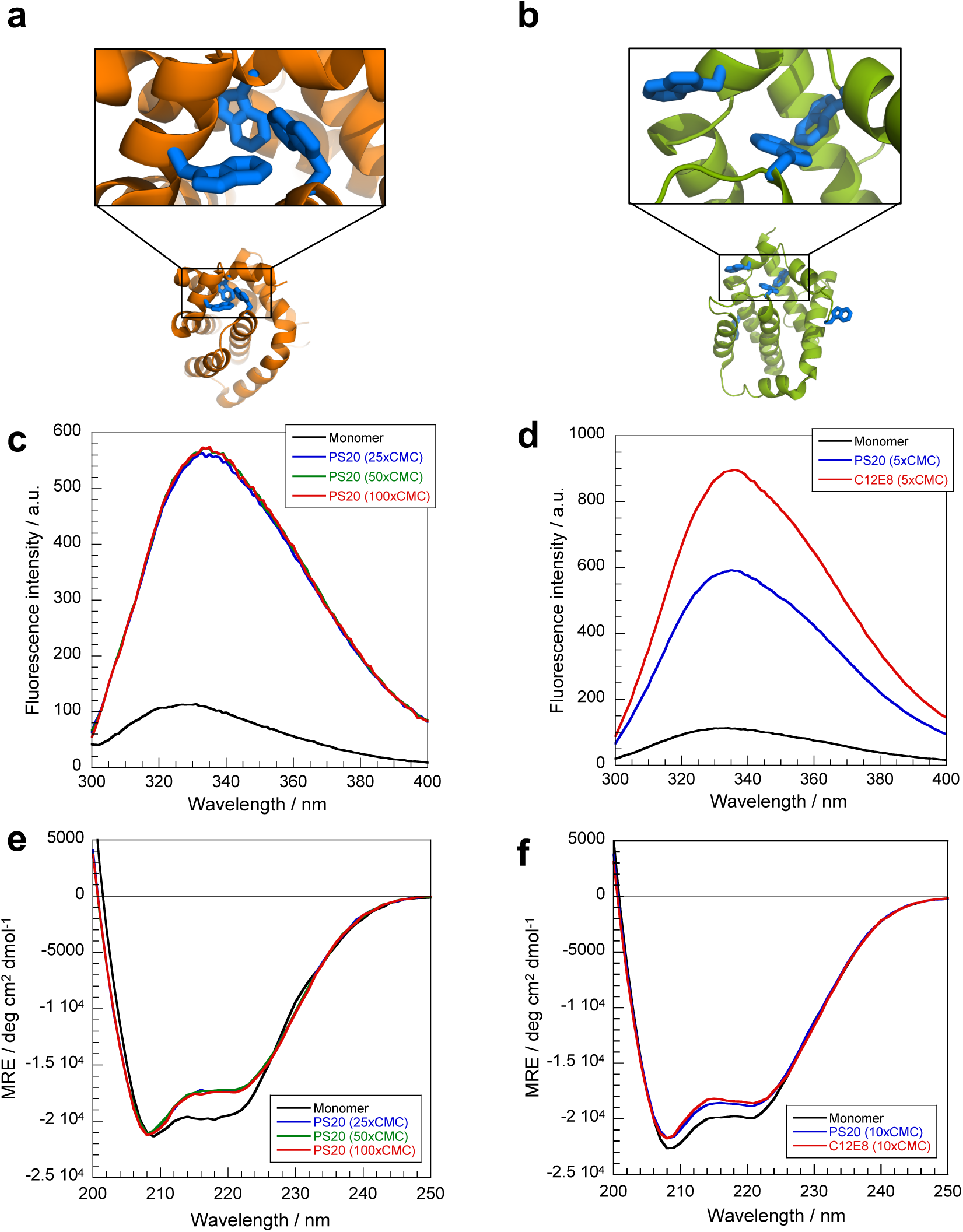
Spectroscopic signatures of BAK (left) and BAX (right) monomers and oligomers. **a,** Structure of BAK (PDB:2YV6). **b,** Structure of BAX (PDB:1F16). Tryptophan residues are shown in blue, and the clusters highlighted. **c, d,** Fluorescence emission spectra of monomers (black lines) and oligomers (colored lines). Excitation wavelengths were 295 and 280 nm for BAK and BAX respectively. **e, f,** Circular dichroism spectra of monomers (black lines) and oligomers (colored lines). Protein concentrations were 5 and 10 μM for BAK and BAX respectively. Detergents concentrations are indicated in the legends as multiples of their CMC’s. All spectra were buffer-subtracted.

**Extended Data Fig. 7.**
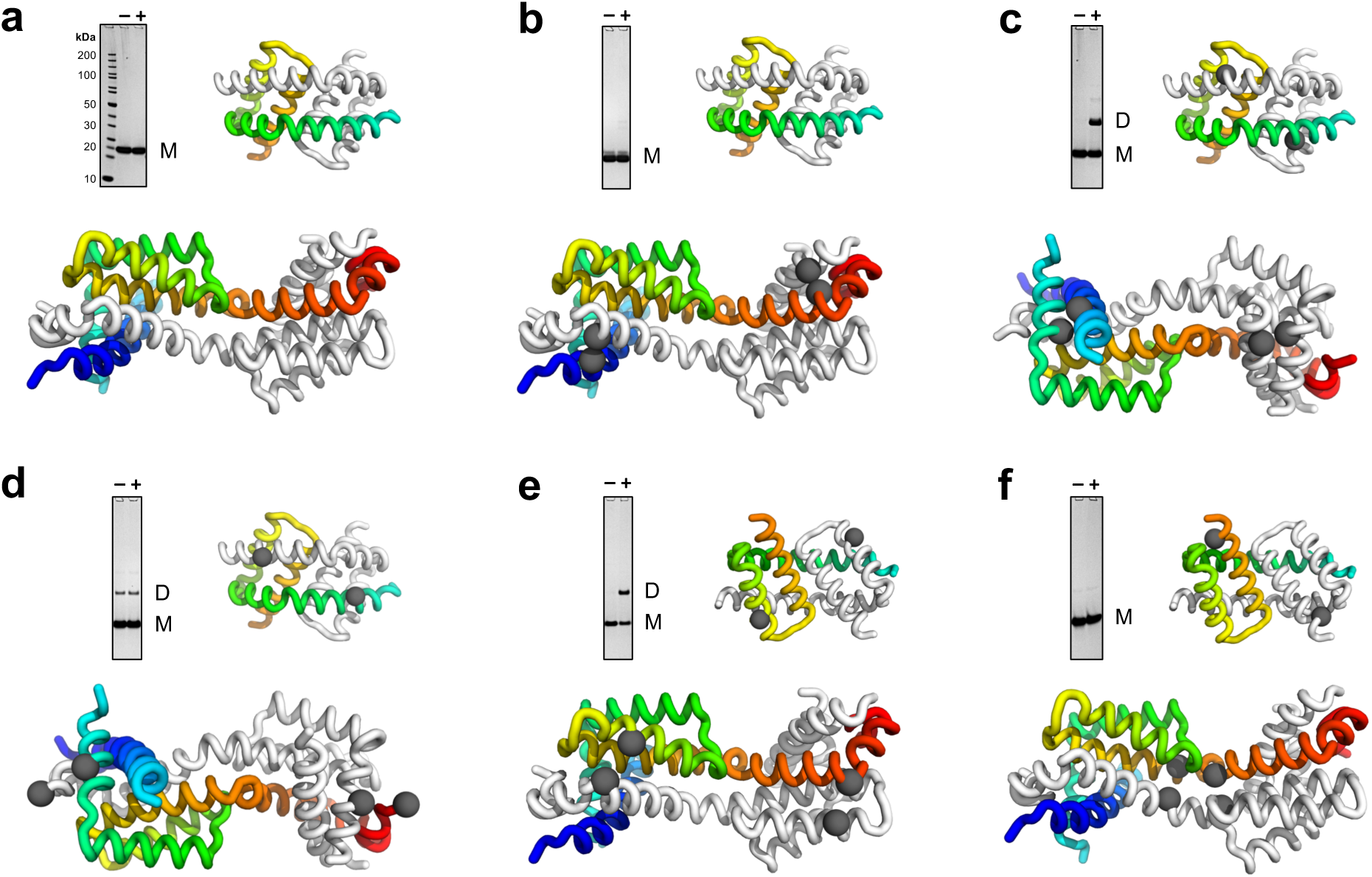
Topological mapping with disulfide staples indicates that the non-physiological helix-swapped dimer of BAK (PDB:4U2U, bottom) is not compatible with the oligomeric structure formed in PS20. We inserted cysteine residues to allow formation of cross-linking ‘staples’. The positions of these mutations are indicated on the structures as grey spheres. Inter-molecular disulfide bond formation (dimerization) was assessed by denaturing SDS-PAGE in the presence (+) or absence (–) of PS20 (20xCMC). **a,** ‘Wild-type’ BAK (no cysteines). **b,** A28C/L163C. **c,** Y41C/A79C. **d,** Q77C/Q184C. **e,** T116C/H165C. **f,** V142C/F150C. Both the absence of dimer formation in (**b**), (**d**) and (**f**), and the presence of disulfide dimers in (**c**) are inconsistent with the helix-swapped dimer structure from Brouwer *et al.* (2014) (PDB:4U2U, bottom). Thus, the oligomers formed in PS20 are different from this non-physiological dimer. Structures are colored from their N-terminus (blue) to their C-terminus (red) for one protomer, and grey for the other one. Note that the data did not allow us to assess the validity of the BH3-in-groove dimer model (Brouwer et al., 2014) (PDB:4U2V, top structure), since this structure only contains helices α2–α5 (highlighted by matching the color gradient to aid comparison).

**Extended Data Fig. 8.**
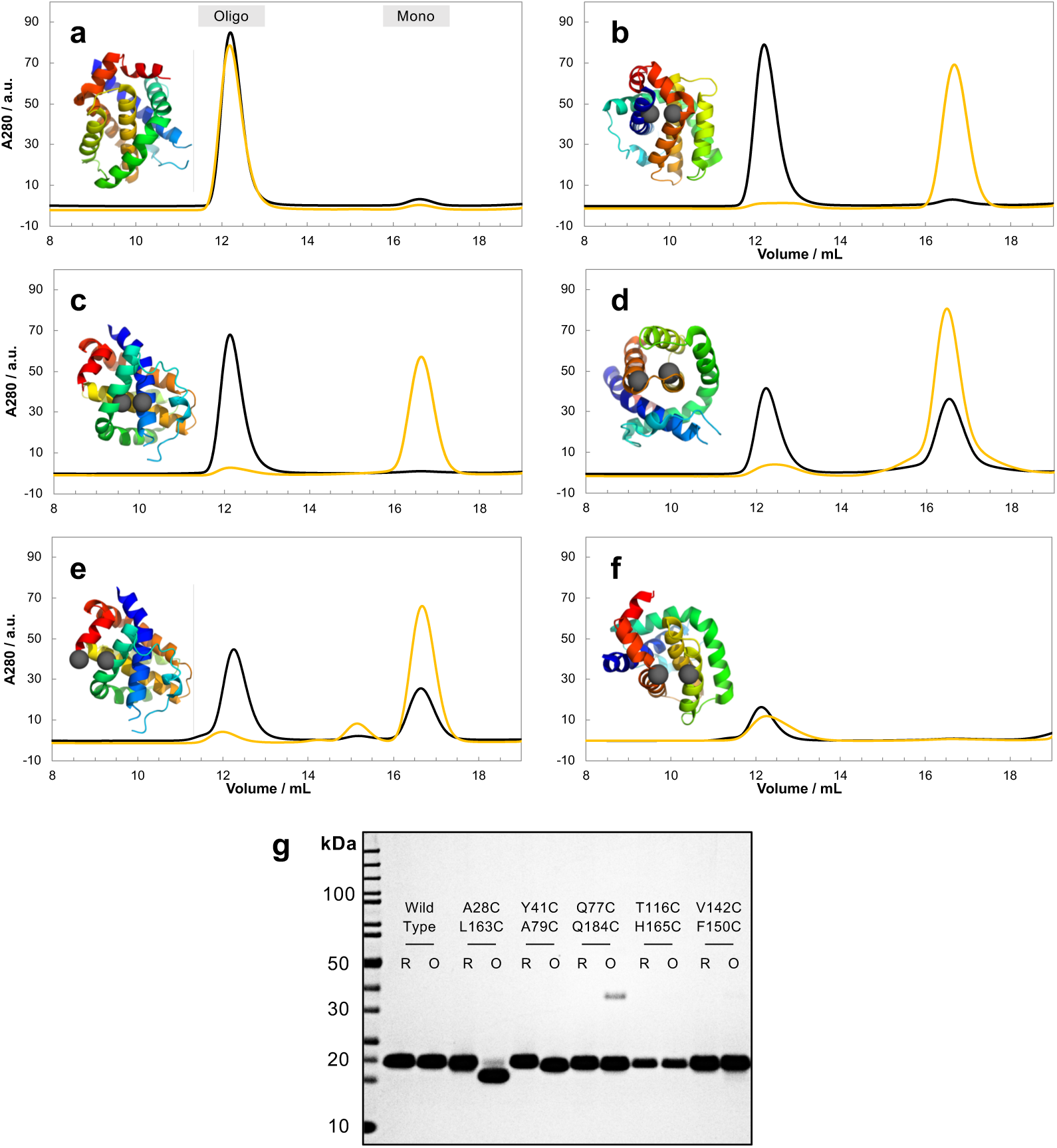
Disulfide mutants of BAK which are incapable of MOMP also suppress oligomerization in detergent. Iyer *et al.* (2016) showed that specific disulfide cross-links (shown in panels **b**, **c**, and **d**) inhibit the ability of BAK to form pores and induce MOMP. The same cross-links suppressed oligomerization in detergent. For each double mutant, 20 μM of protein was treated with PS20 (20·CMC), and the oligomerization assessed by SEC (Superdex 200 10/300). The disulfide form (yellow) was compared to the dithiol form (black) to deconvolute the effect of mutations from stapling. Structures of BAK (PDB:2YV6) with cysteine mutations shown as grey spheres are indicated for each double mutant. **a,** Wild-type BAK (control, no cysteines) oligomerizes fully in detergent. **b,** A28C/L163C. **c,** Y41C/A79C. **d,** V142C/F150C. **e,** Q77C/Q184C. **f,** T116C/H165C (at 6 μM). Note that (**e**) and (**f**) were included as further controls, and have not been assessed in mitochondrial assays. **g**, Denaturing SDS-PAGE of oxidized (O) and reduced (R) forms of the double mutants.

**Extended Data Fig. 9.**
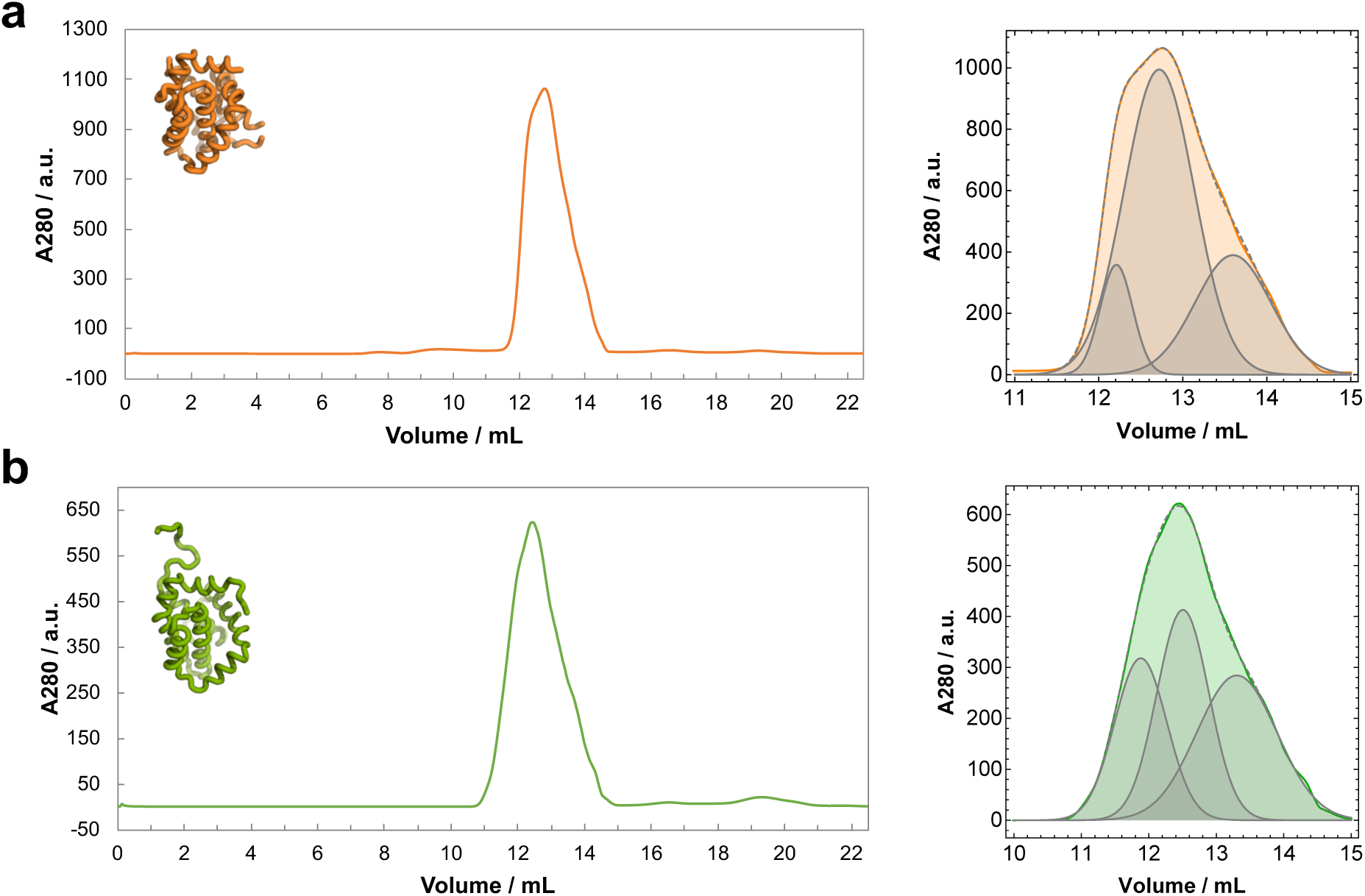
BAK and BAX oligomers appear heterogenous by SEC at high concentrations. **a,** SEC of BAK at ∼450 μM in the presence of ∼90 mM PS20 (1100·CMC). **b,** SEC of BAX at ∼250 μM in the presence of ∼90 mM PS20 (1100·CMC). Right panels show the oligomer peak fitted to a triple Gaussian function (dashed grey line). The individual Gaussian functions are reproduced for visualization (shaded grey). SEC was performed on a Superdex 200 10/300.

**Extended Data Fig. 10.**
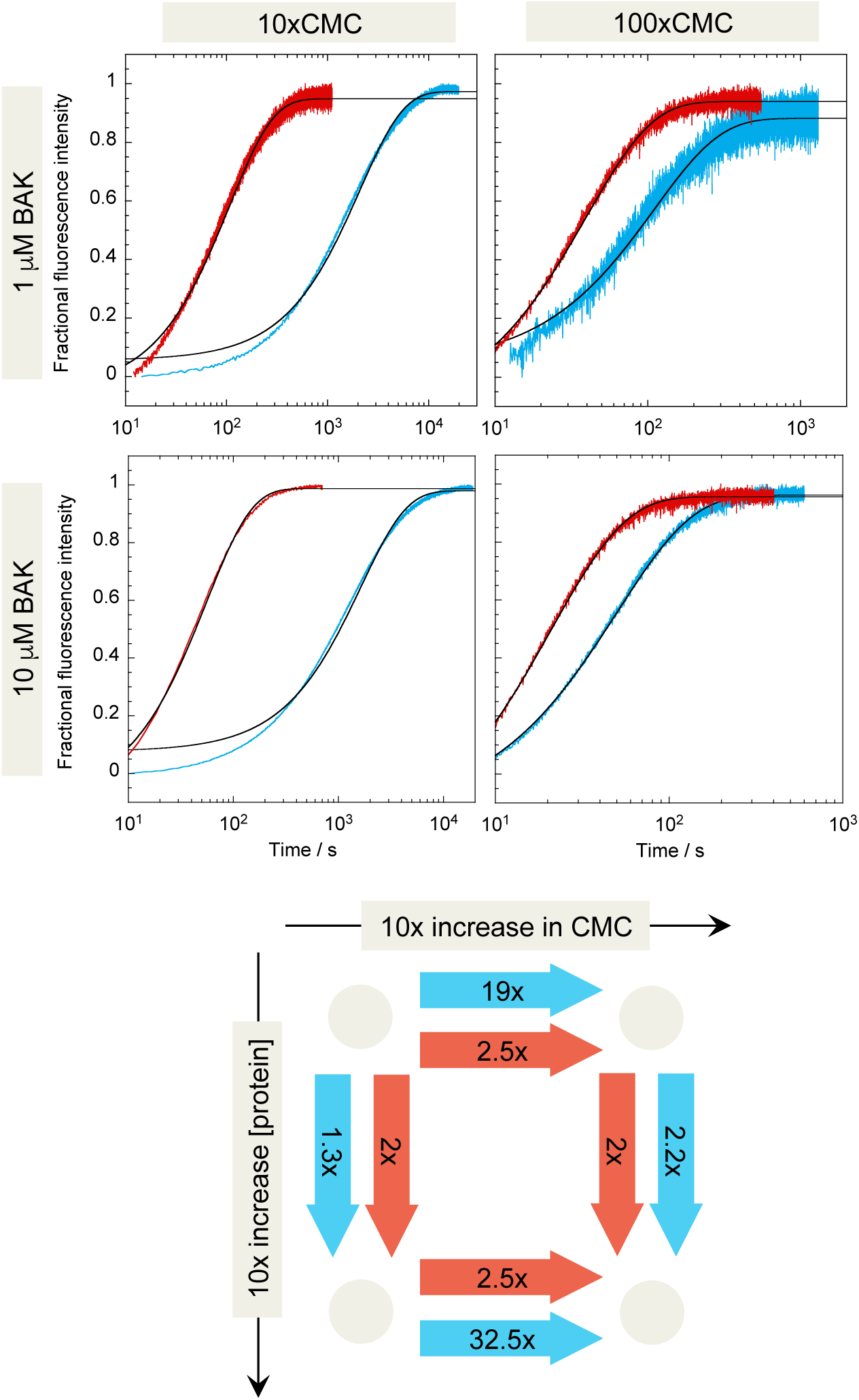
Oligomerization kinetics of BAK as a function of detergent-type. Both C12E8 (red) and PS20 (blue) were tested over a 10-fold range against BAK. The protein concentration was also varied over a 10-fold range. Black lines represent fits to a single exponential decay function. Note the different time-scales. The diagram at the bottom illustrates the approximate fold-change in rate when switching between the protein-detergent conditions indicated by the arrow. A 10-fold change in protein concentration has almost no effect on the rate of oligomerisation. Interestingly, PS20 (blue) appears to have a much greater impact than C12E8 (red) on the rate of oligomerisation.

**Extended Data Fig. 11.**
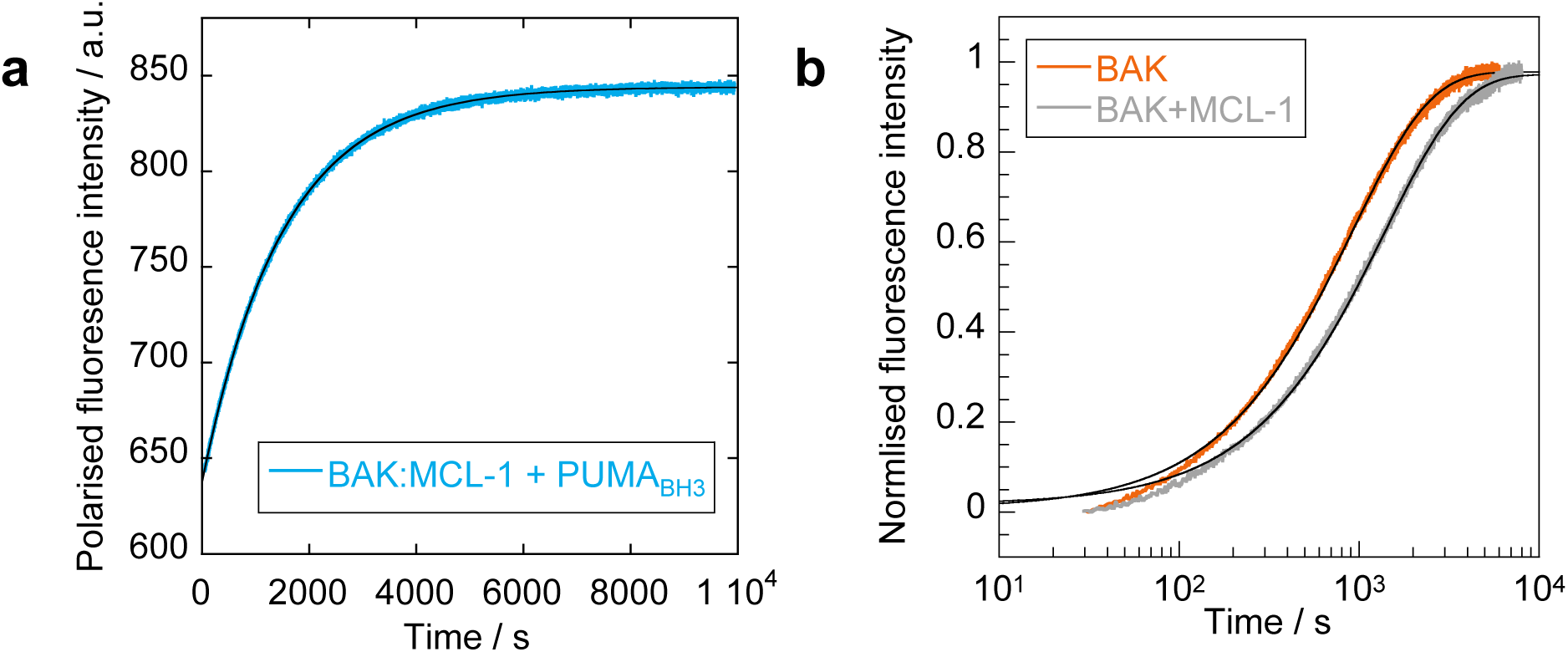
Formation of BAK:MCL-1 hetero-dimers and displacement by PUMA are slow processes. **a,** Binding of PUMA to BAK:MCL-1 hetero-dimers is rate-limited by the dissociation of the complex. BAK and MCL-1 were pre-incubated in the presence of PS20 (20·CMC) to trigger hetero-dimerization, followed by the addition of dye-labeled BH3 peptide. The association reaction was followed by monitoring the change in TAMRA fluorescence polarization, and the data fitted to a single exponential decay function (black line, *k*_obs_ = 6.7 (±0.1) 10^-4^ s^-1^). Binding is extremely slow, and the observed association rate constant is similar to the dissociation rate constant of BAK_BH3_ from MCL-1 in buffer (*k*_off_ = 5.6 (±0.6) 10^-4^ s^-1^), suggesting a dissociation-limited binding event. **b,** The homo-oligomerization of BAK, and its hetero-dimerization with MCL-1 appear to be rate-limited by similar processes. Components were pre-assembled in buffer (5 µM final concentrations), and the reaction initiated by the addition of PS20 (20×CMC). The data were fitted to single exponential decay functions (black lines). Observed rate constants were 1.1 (±0.1) 10^-3^ s^-1^ and 0.7 (±0.1) 10^-3^ s^-1^ for BAK and BAK+MCL-1 respectively.

